# Deliver on Your Own: Disrespectful Maternity Care in rural Kenya

**DOI:** 10.1101/586693

**Authors:** Adelaide M Lusambili, Violet Naanyu, Terrance J. Wade, Lindsay Mossman, Michaela Mantel, Rachel Pell, Angela Ngetich, Kennedy Mulama, Lucy Nyaga, Jerim Obure, Marleen Temmerman

## Abstract

**Background:** Under the Free Maternity Policy (FMP), Kenya has witnessed an increase health facility deliveries rather than home deliveries with Traditional Birth Attendants (TBA) resulting in improved maternal and neonatal outcomes. Despite these gains, maternal and infant mortality and morbidity rates in Kenya remain unacceptably high indicating that more work needs to be done.

**Aim:** Using data from the Access to Quality Care through Extending and Strengthening Health Systems (AQCESS) project’s qualitative gender assessment, this paper examines and describes women’s experience of disrespectful care during pregnancy, labour and delivery. The goal is to promote improved understanding of actual care conditions in order to develop interventions that can lift the standard of care, increase maternity facility use, and improve health outcomes for both women and newborns.

**Methodology:** We conducted sixteen focus group discussions (FGDs) with female adolescents, women, men and community health committee members. Twenty four key informants interviews (KII) including religious leaders, local government representatives, Ministry of Health (MOH) and local women’s organizations were conducted. Data were captured through audio recordings and reflective field notes.

**Research site:** Kisii and Kilifi Counties in Kenya.

**Findings:** Findings show Nursing and medical care was sometimes disrespectful, humiliatings, uncompassionate, and neglectful. In both sites, male health workers were the most preferred by women as they were friendly and sensitive. Young women were more likely to be abused and women with disabled children were stigmatized.

**Conclusions:** Kenya needs to enforce the implementation of the quality of care guidelines for pregnancy and delivery, including respectful maternity care of pregnant women. To make sure these procedures are enforced, measurable benchmarks for maternity care need to be established, and hospitals need to be regularly monitored to make sure they are achieved. Quality of care and compassionate and caring staff may lead to successful and sustainable use of facility care.

## INTRODUCTION

Few health interventions have greater potential impact on the overall health of society than good quality facility-based, care to women while pregnant and during and after childbirth. In Kenya, under the Free Maternity Policy (FMP), more women have been choosing to give birth in maternity care facilities rather than at home with traditional birth attendants (TBA) (1,2). Despite these gains, infant mortality and morbidity rates in Kenya remain unacceptably high and anecdotal evidence shows that not all pregnant women may be willing to attend facility care services. One of the factors that has been shown to affect utilization of facility-based, maternal health care services is the experience of disrespectful care received by women. In the maternity health services context, ‘respectful care entails respect for beliefs, traditions and culture, and [the] empowerment of the woman and her family to become active participants in health care. Respectful care also encompasses continuity of care, the right to information and privacy, good communication between client and provider, and use of evidence-based practices’ (3–5).

A wealth of evidence confirms that women who perceive receiving substandard and disrespectful care during a childbirth are far less likely to seek skilled birth care during subsequent pregnancies (6–8). In addition to increasing their own and their newborns’ risk for poor outcomes, women with these experiences may also discourage others from seeking facility-based care (9).

Research evidence indicates that gender inequalities and unequal power distribution may act as a barrier to respectful maternity care (10,11). Women from low-resource poor settings are likely to experience gender inequalities and to be discriminated against by service providers due to their low status in the society (11,12). Further evidence illustrates that the unequal distribution of power between men and women, and women’s lack of autonomy in decision making at the household levels in these low resource settings exposes them to long term health risks – with enormous social and economic implications (13). Over the past two decades, the Respectful Maternity Care movement has gained considerable attention worldwide as a basic right of women throughout pregnancy, labour, and delivery. The World Health Organization (WHO) further recognizes the principle of respectful care as a major factor for increasing the use of pregnancy and maternity healthcare services, and ultimately, better maternal and neonatal outcomes (3,4,14).

Even in HICs, where there are professional standards and documented policies with respect to delivery of care, legal consequences for violations in providing care, and patient familiarity with their rights when receiving care, literature illustrates that vulnerable women in low social economic status may experience disrespectful care (15).

Research evidence from low resource settings has reported some incidents of individual behaviour and practices of caregivers at health facilities who violate the principles of respectful and compassionate maternity care (16–19). In part, this research points to a lack of adequate staffing, outdated equipment, and a lack of enforcement of explicit standards for professional ethics as a cause of disrespectful care.

Lacking adequate basic components to ensure the delivery of respectful and suitable health care, it is easy for health care workers to engage in substandard and disrespectful care such as routinely ignoring patient requests. In fact, evidence from some LMICs demonstrate that women about to give birth, especially poor and rural women, are more likely to be neglected, humiliated, and often subjected to verbal and, at times, physical abuse (16,18,19). In Kenya, anecdotal evidence indicates that some pregnant women seeking maternity care are likely to be abused, however, there is little research evidence that corroborates these views.

In an attempt to begin to address this gap in the literature, this study analyzed data from a qualitative gender assessment of the Access to Quality Care through Extending and Strengthening Health Systems (AQCESS) project to describe and shed new light on how Kenyan women in two dissimilar rural settings experienced maternity care. The project was executed by Aga Khan Foundation Canada (AKFC) and implemented by agencies of the Aga Khan Development Network (AKDN) with financial support from the Government of Canada, through Global Affairs Canada (GAC) and the Aga Khan Foundation Canada (AKFC).

## METHODS

Qualitative in design, the overall aim of the study was to provide preliminary evidence to begin to understand the gendered dimensions of access to and control over resources, decision-making, social norms and perceptions and practices related to access and use of MNCH services in LMIC countries.

The data come from two target communities -in Kenya’s rural Kisii and Kilifi Counties. Kisii area is located in south-western Kenya and Kilifi is a historical coastal county located northeast of Mombasa. These two locations were chosen because they are the target locations for the AQCESS project. The intention was to inform the AQCESS project implementation in these two locations to ensure gender responsiveness of interventions.

**This paper aims to build on the learnings of that assessment and focus in more** detail on the experience of some participants related to disrespectful maternity care. Seeking to elicit representative views, the researchers conducted 24 Key Informant Interviews (KIIs) with a wide variety of stakeholders including health workers, government and religious leaders, women representatives, and service providers. In addition, 16 Focus Group Discussions (FGDs) were conducted across the two research sites, eight (8) in each. To allow full and free participation, two gender responsive FGDs from each category, - adult males, adult females, adolescent females, and members of the community health committees were interviewed separately.

Based on the mounting evidence that distance plays a central role in gaining access to health care (20,21), focus group participants and interviewees were selected from sites wherehalf of each FGDs were located ≤5 km long distance and half were located ≥ 5 km from the nearest maternity health facility.

The Aga Khan University’s (AKU’s) Ethics Review Board and the National Commission for Science, Technology, and Innovation (NACOSTI) approved the study. In addition, approval to commence data collection was also approved by the County offices at both sites. Data in both sites were collected in April 2017. Before participating, all respondents gave informed consent. Researchers explained the purpose of the study, the potential risks, and that participation was voluntary and participants could withdraw at any time. Community Health Volunteers (CHV) oversaw and confirmed the consent of participants younger than 18 years old, who also provided their individual assent to participate.

Qualified moderators conducted FGDs in Swahili, local dialects, or English, as appropriate. All discussions and Interviews took no more than 2 hours. The field team received training on the protocol, tools, and on gender sensitive research and ethics. Interview guides for both participants and interviewers were available in relevant languages. During the interviews, all sessions were recorded with permission of respondents. Data needing translation from Swahili or local dialects to English were translated. Transcribed data were coded, encrypted, and saved securely in accordance with the AKU’s Ethics and Data Protection Act.

## DATA ANALYSIS

Upon the completion of the fieldwork in 2017, qualitative data were analyzed using a continuous iterative process (22). Each transcript was coded independently by two analysts and reviewed by one study investigator who was the chief data analyst. Where coding discrepancies occurred, at least two analysts re-examined the transcripts and discussed all possible meanings associated with the text in question until agreement was achieved. This paper aims to build on the learnings of that assessment and focus in more detail on the experience of participants related to disrespectful maternity care. An additional coding of key themes is illustrated in table 1 below. The current analysis relied on the users’ own report of their experience of disrespectful maternity care services.

**Table 1:**
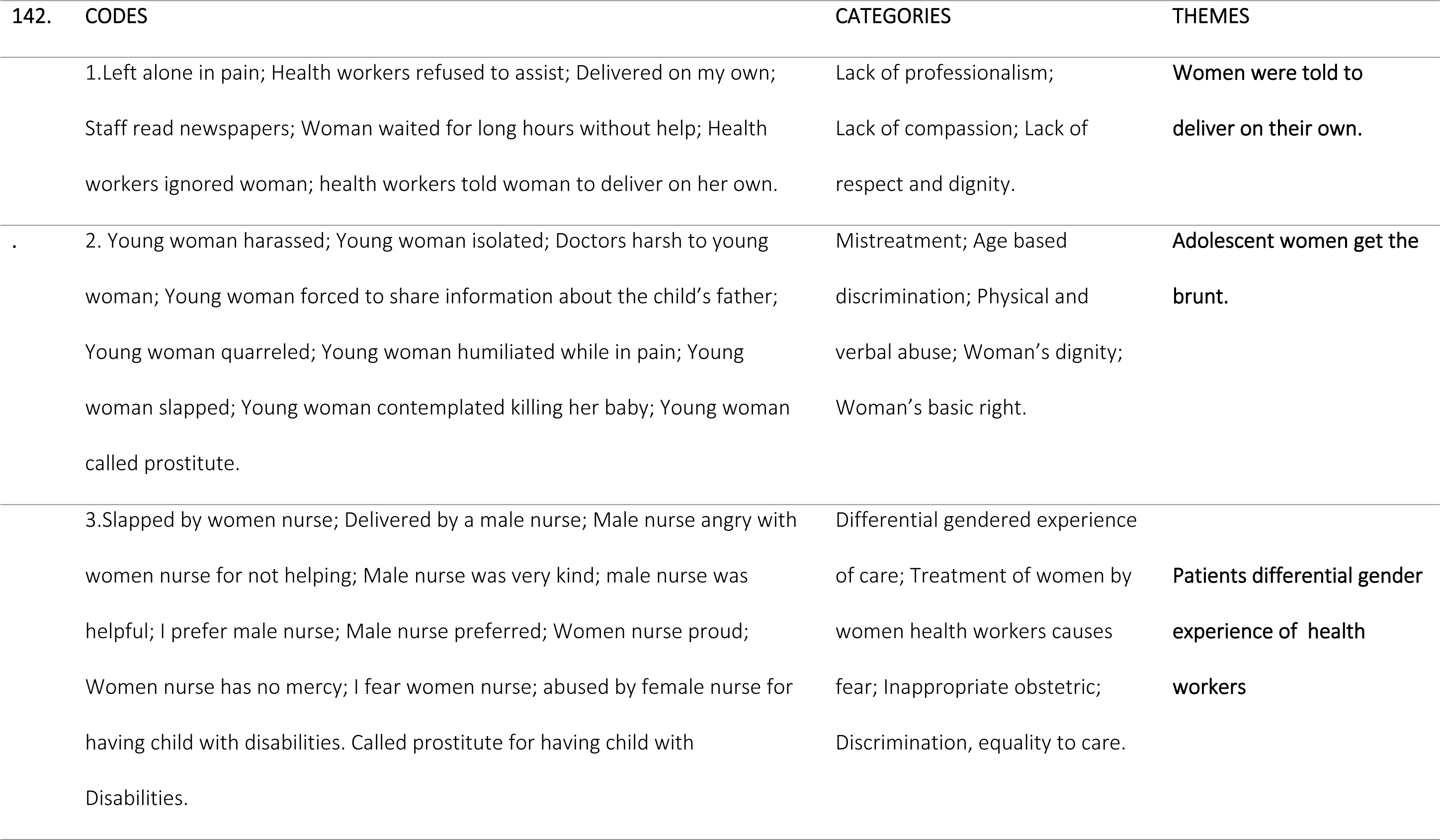
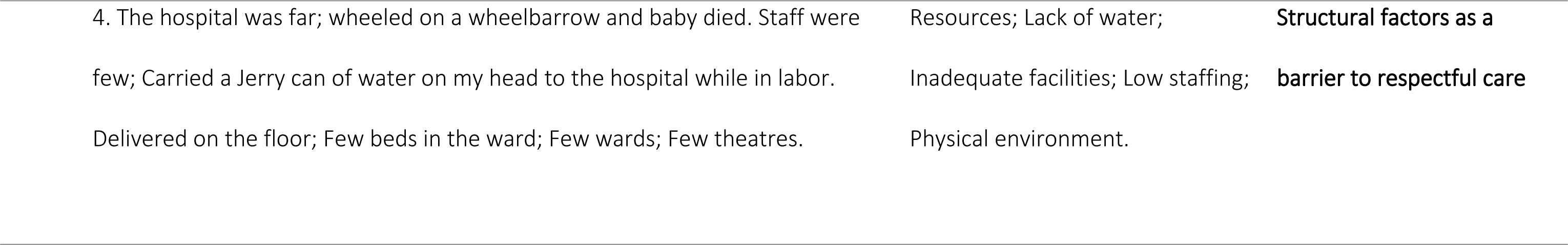
Codes, Categories and Themes

| 142. | CODES | CATEGORIES | THEMES |
| --- | --- | --- | --- |
|  | 1.Left alone in pain; Health workers refused to assist; Delivered on my own;<br>Staff read newspapers; Woman waited for long hours without help; Health workers ignored woman; health workers told woman to deliver on her own. | Lack of professionalism;<br>Lack of compassion; Lack of respect and dignity. | **Women were told to deliver on their own.** |
| . | 2. Young woman harassed; Young woman isolated; Doctors harsh to young woman; Young woman forced to share information about the child's father; Young woman quarreled; Young woman humiliated while in pain; Young woman slapped; Young woman contemplated killing her baby; Young woman called prostitute. | Mistreatment; Age based discrimination; Physical and verbal abuse; Woman's dignity; Woman's basic right. | **Adolescent women get the brunt.** |
|  | 3.Slapped by women nurse; Delivered by a male nurse; Male nurse angry with women nurse for not helping; Male nurse was very kind; male nurse was helpful; I prefer male nurse; Male nurse preferred; Women nurse proud; Women nurse has no mercy; I fear women nurse; abused by female nurse for having child with disabilities. Called prostitute for having child with Disabilities. | Differential gendered experience of care; Treatment of women by women health workers causes fear; Inappropriate obstetric; Discrimination, equality to care. | **Patients differential gender experience of health workers** |

|  |  |  |
| --- | --- | --- |
| 4. The hospital was far; wheeled on a wheelbarrow and baby died. Staff were few; Carried a Jerry can of water on my head to the hospital while in labor. Delivered on the floor; Few beds in the ward; Few wards; Few theatres. | Resources; Lack of water; Inadequate facilities; Low staffing; Physical environment. | **Structural factors as a barrier to respectful care** |

## FINDINGS

Pregnant women seeking antenatal care as well as women in labour during and after delivery report that some health care workers in Maternity Care Facilities are at times disrespectful, demeaning, humiliating, verbally or physically abusive, dismissive, and neglectful of patient-reported pain, and suffering. Although our results illustrate that women of all ages experienced these types of inappropriate care and neglect and abuse, younger women and especially adolescent women, were more likely to report verbal and physical abuse. As well, both male and women focus group participants reported that female health workers were more likely to demonstrate abusive behaviors and, as a result, many participants indicated that they preferred to be tended to by male health workers. Below, we present the specific themes that emerged from the data analysis and provide vignettes to illustrate and support them.

*The following themes emerged from the data.*

1. Women were told to deliver on their own
2. Adolescent women get the brunt
3. Patients differential gender experience of health workers
4. Structural factors a barrier to respectful maternity care

### 1. Women were told to deliver on their own

Data from both research sites confirmed cases in which patients’ basic right to care was violated. Speaking of her own personal experience with maternity healthcare workers, a participant one of the women FGDs reported that some of the health workers refused timely help to women at the time of delivery.

> …My experience with my first-born delivery was not good with a women nurse. I told. her I was in pain, and she abused me saying I should ‘stop my nonsense and wait to give. birth in the morning.’ Soon after, a male nurse came and assisted me [to] give birth. He even quarreled [with the women nurse], asking her why she was treating me that way. (Women FGD, Kisii).

In the following, a mother described her experience while accompanying her relative, and the treatment her cousin received convinced her to never again go back to a public health care facility.

> Haaa, I can’t, I better go to the private clinic … With those mockery and abuses I can’t. 1Like when we went at night with my cousin. There was a male doctor who we think was 1on drugs—chewing miraa, was sitting outside, maybe he was taking bhang. He told the 1girl ‘it’s not you [who can] tell us what to do!’ When the pain was too much, she went to the bed on her own, and the man came wore glove inserted his fingers and literally tore the lady… (Adolescent Women FGD, Kisii).

Key informant interview with a local politician highlights the pattern of neglectful and uncompassionate approach to care at the facility.

> …I have witnessed a case at the hospital where there was a woman who came for the first. time and she was told she would not be attended to until she brings her husband and get. tested, she went and never came back (Women representative KII-Kisii).

In the following example, a participant in an adult male FGD speaks of his experience with staff while accompanying his relative to the hospital.

> I took my sister in-law to hospital, and after assessing her, they left her alone in bed. She called for assistance, but they said, ‘It’s not yet time.’ She pushed and delivered on her own, that’s when the health workers came running wiping the child…at times they can beat you because you gave birth on the floor, as if the mistake is yours. Yet they don’t respond when called (Male FGD, Kisii).

Some healthcare staff who neglected patients were reading newspapers. Healthcare workers sometimes left the facility altogether, going for long lunch “hours” and leaving patients for hours on end, waiting to be attended to:

> I can add that medical providers are very… abusive especially for young ladies and adolescents. Some medical staff decide to go for early lunch. Some read newspapers 208. while patients are waiting for services (Male FGD, Kisii).
>
> It happened that since she declined going to theatre the health workers ignored her and told her to deliver on her own, but God intervened [by chance]. So the nurse’s attitude, Ignorance could be one of the things that contributes to this. (Male FGD, Kisii)

The above vignettes demonstrate some of the ways women were treated while under the care of service providers. The vignettes also demonstrate a lack of professionalism and compassion among some health workers.

### 2. Adolescent women get the brunt

While some of the adult women reported that they experienced different kinds of incivility and poor care, the adolescent women focus groups reported that healthcare workers in both Kisii and Kilifi were more likely to abuse very young women, both verbally and physically. A focus group discussion among adolescent girls in Kisii confirmed that some doctors also share this prejudice. In this case, a doctor uses harsh and humiliating language when speaking to young pregnant women during the delivery process.

> … During the pain, they [doctors] abuse you and tell you to deliver. Yet you don’t know anything. They tell you to push, and you don’t know how.” (Women Adolescents FGD, Kisii).
>
> …We (adolescents) are attended to, but harshly. For example, me, I’m young and 225. they told me, ‘Young as you are, you went opening your legs to men.’ (Women Adolescents FGD, Kisii)

Some adolescent women reported being slapped during the labour process. One was so upset by abusive nature of some staff, she contemplated killing the child.

> she [the nurse] was slapping me. I said, ‘I’ll kill the baby and give birth to a dead baby,’ but the lady. who escorted me went and called for a male doctor, who assisted me… (Women Adolescents FGD, Kilifi)
>
> … Some doctors are harsh because you’re a girl, during the pain they abuse you and tell you…’Young girl, when you were loitering looking for that pregnancy, were we there?’Imagine! Instead of helping you… (Women Adolescents FGD, Kisii).
>
> … They ask you questions, ‘When did you get pregnant? Who gave you this pregnancy?’ You don’t know what to answer them. If you don’t answer, the leave you there unattended, and say ‘Till you answer …’ (Women Adolescents FGD, Kisii).

As the following discussion from the male FGDs illustrates, this kind of poor treatment appears to stem from social disapproval, leading to disrespect for adolescent girls who become pregnant.

> *Respondent:* Our young wives, usually they encounter a lot of problems.
>
> *Moderator:* Oh, young wives? Ha!
>
> *Respondent:* Yes, the young ones. You know us. We have been, ah, so
>
> (laughter) … when they go there, and especially because they are young, they complain that they are harassed and asked: ‘You! As young as you are you …,’ [and] such like harassment.
>
> Moderator: Why such harassment?
>
> *Respondent:* Because they are young and are already pregnant. And what are they expected to do, and it has already happened?
>
> *Moderator:* Okay. So how is she harassed? (Group reaction)
>
> *Respondent:* She gets uncomfortable.
>
> *Respondent:* They begin to think ‘this young girl has already conceived,’ so when she goes there to be served, she feels out of place and fearful, hence feel she has not been served well. She feels like she has been isolated (Male FGD, Kilifi).

While some women generally suffer abuse from maternity facility caregivers, levels of physical abuse were reported to be greater for younger women during the actual delivery.

> *Respondent:* Actually, it is during delivery that there are usually a lot of difficulties.
>
> *Moderator:* Kindly mention all.
>
> *Respondent:* Some of [the women] are slapped, and that story became a talk of the town some time back, yeah. Some of the doctors [show] that behavior. (Okay). ‘You did it willingly, and you want to cause us trouble now.’ You see, that’s bad to tell somebody. She might decline to come back again, when she is pregnant in the future, and will prefer to home delivery. … Yeah, there is such a one [who slaps the women] here, only that I cannot disclose, but they are there. Very short-tempered, even tells you not to come back to that hospital (Male FGD, Kisii).

Poor treatment of young women was also reported in some of the Key Informant Interviews. In the following vignettes, two Ministry of Health (MOH) representatives observe:

> …the way they [staff] handle these mothers, somebody may harass the mothers and next time… or even when she goes back she will go with a bad picture and says,’ I cannot go back to that facility, they do not handle people properly, they call us with very abusive words …. (MOH representative KII –Kisii).
>
> …“I guess once again I would say the adolescents are quite disadvantaged because even the stigma is within the medical care providers, so in terms of the MCH, it is very difficult for the adolescent mothers because they need to seek health care, yet providers’ attitude is the worst when dealing with adolescent mothers.” (MoH representative -Kisii)

These vignettes illustrate the differential experience of maternity care among adolescent mothers in two settings. Moreover, the similarity among the vignettes from both the FGDs and key informants show a consensus regarding the greater level of disrespectful care that these adolescent mothers must tolerate.

### 3. Patients differential gender experience of health workers

The data showed that some women in Kenyan maternity care facilities generally found that female health care workers were more likely to be disrespectful and abusive compared to male care workers. Some participants in the focus groups expressed their preference for male health workers, who they indicated treated them better and were more willing to help when asked. Young women, as a group, were the most vocal in expressing their opinion of female staff and their preference for male health care workers.

> …especially ladies, they are so harsh; they think they dropped from heaven [special than everyone] and us were collected. Men are better; they can tell you to push while assisting you. But females will slap you, yelling at you to open your legs while shouting ‘I want to see the child!’ with abuses. She tells [me] ‘I am waiting to hold the child! (Female Adolescebt FGD, Kisii).
>
> … Especially female doctors are the worst. Men don’t have problems. If you come with a Range Rover [or a Mercedes] Benz, they will wheel you to the ward. But if brought with a wheelbarrow, you’ll be told to move from here to there—they don’t mind the pain you have. You’ll be locked inside a room, and be told to yell there. I was locked [in] and told go for long call there (Female Adolescents FGD, Kisii).

While participants interviewed for this study preferred male attendants as they provided more respectful care, Muslim patient’s desire for male caregivers was tempered by cultural and religious beliefs. A Muslim participant in the adult male focus group reported how his wife resisted being assisted by a male worker.

> I think the most interesting thing is that the women prefer to be served by male health workers rather that the females…Yes that is the truth but not so for us as Islamic. My wife declined totally to be assisted by a male nurse. Even during the clinic, she forbids the male nurse to even touch her when she is to be injected…Actually, it happened that the baby’s head was already out before she could even be assisted. The only person that was available was a male nurse. You know in our religion we prefer not to show the nakedness of a woman - it is sacred… (Male FGD, Kilifi).

Female health workers were reported as being more likely to be verbally and physically abusive. As a result, patients therefore generally preferred to be attended by male health workers.

> *Moderator:* So, you are saying that women prefer the male health workers to the female?
>
> *Respondents:* Yes! (Chorus answer)
>
> *Respondent:* Yes, because the male nurse has a patient heart. He will even try to console the mother, unlike the female nurse, who can inflict slaps. Then she goes about her businesses, leaving the mother behind without even caring (Women FGD, Kisii).

Participants reported that women who gave birth to children with disabilities were likely to be humiliated by some female staff who blamed the mother for the child’s disability.

> They are women and maybe they’ve also given birth. They should know the pain they went through … If you give birth to a disabled child, they ask you when coming to the clinic, ‘Were you moving with men when pregnant?’ Or, ‘Your man did fix you well’ (hakuingisha [penetrate] vizuri [good or satisfactory]) so the child didn’t reproduce properly. The man didn’t have energy.’ (Female Adolescents FGD, Kisii).
>
> …female health workers have contempt. When we went with my cousin, who is also young, we brought a disabled child. Another lady nurse abused us, till we also abused her back. We looked for another doctor, the nurse abused her saying: ‘She was sleeping with men while pregnant, that’s why she gave birth to a disabled child!,’ and we reported the case to the senior doctor…We don’t know if she was reprimanded. My cousin wanted to kill the child, saying she is abused because of her child status. We later took the child to my grandmum, who took care of the child (Female Adolescents FGD, Kisii).

These vignettes illustrate some of the gendered experience of health workers by service users. The vignettes also illustrates some of the attitudes and beliefs associated with giving birth to a child with disabilities, an area that has not been examined in both research and policy in the Kenyan context.

### 4. Structural factors a barrier to maternity respectful care

Our analysis confirmed that, in both Kisii and Kilifi, public prenatal and maternity health care failed to treat some women with dignity and respect. Respectful care is a process in that women must have available structures to support them on their journey through maternity. Findings from this study reveal that some maternity health centres may lack facilities such as water, beds or readily fueled vehicles and ambulances to transport patients that are referred to larger facilities. Participants reported that in some cases women carried water along with them to the hospitals during delivery.

> …You can find a woman who is in labour carrying a jerrican of water on her head going to the hospital. Simply because she knows there is no water at the hospital… (Male FGD, Kilifi).
>
> ..I saw a woman groaning in pain she had come to deliver and there was no water. Usually there is scarcity of water in this area. It happened that that day the care provider present was not supposed to be on duty that night. So it happened as I was talking with her that is when that mother came in but she had to be send elsewhere because the hospital was not functioning to the lack of water… (Male FGD, Kilifi).

Spaces and beds were not adequate to meet the demand of pregnant women seeking care at some facilities.

> The beds in the labour ward should be added. The wards are also small. Some women are usually waiting to give birth while lying down on the floor because the beds are occupied…When I was delivering, I gave birth while lying on the floor because the beds were occupied and there was nowhere to deliver. I knelt and the baby came… (Women FGD, Kilifi).

The available spaces were not gender inclusive and thus discouraging uptake of services.

For some women, structural barriers such as poor road systems and inadequate means of transport hindered accessing the facilities. The male FGDs, both in Kisii and Kilifi observed the difficulties some women face in navigating social structures that are unsupportive and the deleterious impact this had on maternal outcomes.

> …I come from a place called Ibencho, roads are in bad condition. There is a woman who wanted to deliver and was carried using a bed and because of the distance, she died before reaching the hospital…this was less than five years ago. From Ibencho, people are only carried using beds or wheelbarrows to Sengera…Things are not different with Riokindo.” (Male FGD, Kisii)
>
> … I have always witnessed women suffering and having a rough time in accessing the facilities due to long distances that they have to cover. And if it is a must they get to hospital the only available means of transport is the motor bikes. So you can imagine a pregnant mother being rode on a motorbike, it is usually a hard task. This is a challenge. So that is what I have been able to witness also sometimes it happens that some due to that they end up having complications and some even may die before getting to the health facility. This I have witnessed many times and secondly, when they get to the hospital you find that midwives are not available (Male FGD, Kilifi)

These vignettes demonstrate that on a woman’s journey through pregnancy and delivery, there are many barriers that she must navigate both at the micro and the macro levels. Disrespectful care cannot only be considered from the way health workers treat women but must consider additional factors such the availability of resources required to provide respectful and appropriate care.

## DISCUSSIONS AND POLICY IMPLICATIONS

In this analysis, women in both research sites reported mistreatment and lack of respect by some health care workers, and particularly by female staff. These reports, coming both from people who have experienced the public maternity care facilities in geographically separate rural contexts of different sizes as well as those key informants who work in or oversee these facilities suggest that there is a tendency for disrespectful care of pregnant women and women giving birth. Moreover, this deficit in appropriate care appears to be even greater for women who are poor, young, or have children with birth defects, potentially undermining the efficacy and reputation of the entire Kenyan public maternity health care system.

Our findings raise issues around various aspects of delivering acceptable and respectful care including social cultural norms with regard to gendered nature of maternity care, the stigma around age, pregnancy and disabilities, and structural barriers and inadequacies of resources for maternal care. As a result, there is an urgent need to address these various issues to ensure that the Kenyan Free Maternal Policy to provide safe, satisfactory care to reduce infant and maternal mortality is realized. There is also need for a training on cultural change in norms and attitudes that are associated with age and disability across the health system structures. To combat negative attitudes and behaviors, existing standards of care must be enforced. Maternity care facility staff must be supported to understand and deliver these standards of care, with their implementation of these standards consistently monitored.

The consistency of these reports, especially when it comes to the treatment of very young women, shows that these are not isolated cases as they are consistent with research from other low and middle income countries (16–19). Rather than looking at the disrespectful and neglectful nature of some health workers in isolation, a systems perspective theoretical approach can provide insights on how to address some of the mishaps. A system perspective approach posits that behaviors are part of a larger system including all the structures that support that system [in this case: policy makers, leaders, medical boards, national governments, local governments, international regulators [WHO), the patient, service providers, environments etc]. Therefore, to ensure that women receive timely and respectful health care at all times, all health system actors must be engaged to promote an equilibrium functional environment where acceptable behavior in provision of care is monitored and sustained. For example, the World Health Organisation (WHO) guidelines on ‘Care During Pregnancy’ prohibits the routine shaving of hair as it increases infections. How do we ensure that women are not subjected to this without consenting? And do the health workers know that these guidelines exist?. Facilities should employ more midwives –for example, in Kilifi, the nurse to patient ratio is 3 midwives per 10,000 population compared to 23 per 1000 population recommended by WHO. Increase in staffing should be accompanied by incentives to staff, a training package on respectful care and a locum policy.

While it is our responsibility to strive to promote respectful maternity care, we acknowledge that placing the blame solely on health care workers in isolation will not solve the inadequate services pregnant women receive. There is a need for a systemic and institutionalized effort spanning from policies to address community-based, socio-cultural norms, health care educational training, and documented professional associations standards of care and enforcement of these standards. Instituting standards of care in Maternity Care Facilities; re-educating health care workers at all levels; and instituting monitoring plans to make sure that pregnant women, women, and newborns are treated with dignity and respect while receiving obstetrical and neonatal care including coaching and mentorship is critical for shifting attitudes and behaviors.

The fact that female health workers were reported as being more likely than male health workers to abuse women under their care requires further study to determine the underlying factors for these attitudes and behaviour, and measures to address this issue. Maternity Care Facilities will need to monitor all staff closely to make sure patients are treated with kindness, respect, consideration, and professionalism. To reach that goal, staff attitudes towards patients, and the way they treat them needs to be a key element in both hiring and retention, and in the most egregious cases, abusers need to be reported to the police for redress under the law. A structure for reporting and response must be devised and instituted to make sure that, when dealing with patients, staff understand and carry out the principles of Respectful Care and gender responsiveness.

Lastly, the findings raise issues as to whether sufficient training and professional development for health service providers in Kenya is delivered, particularly around respectful maternity care and gender responsive services. There may be a need to review the current curriculum and identify potential areas for interventions.

**Table 2:**
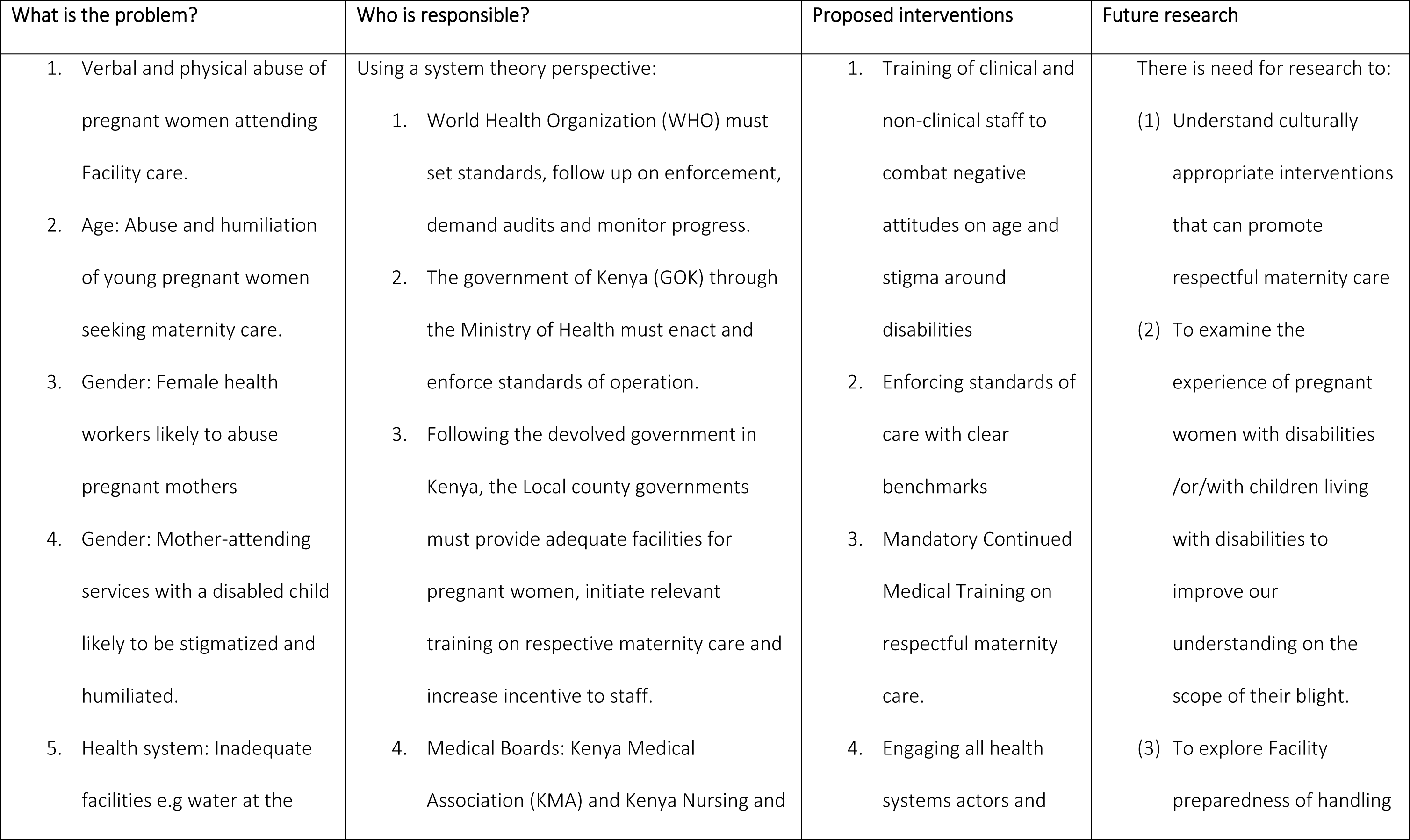
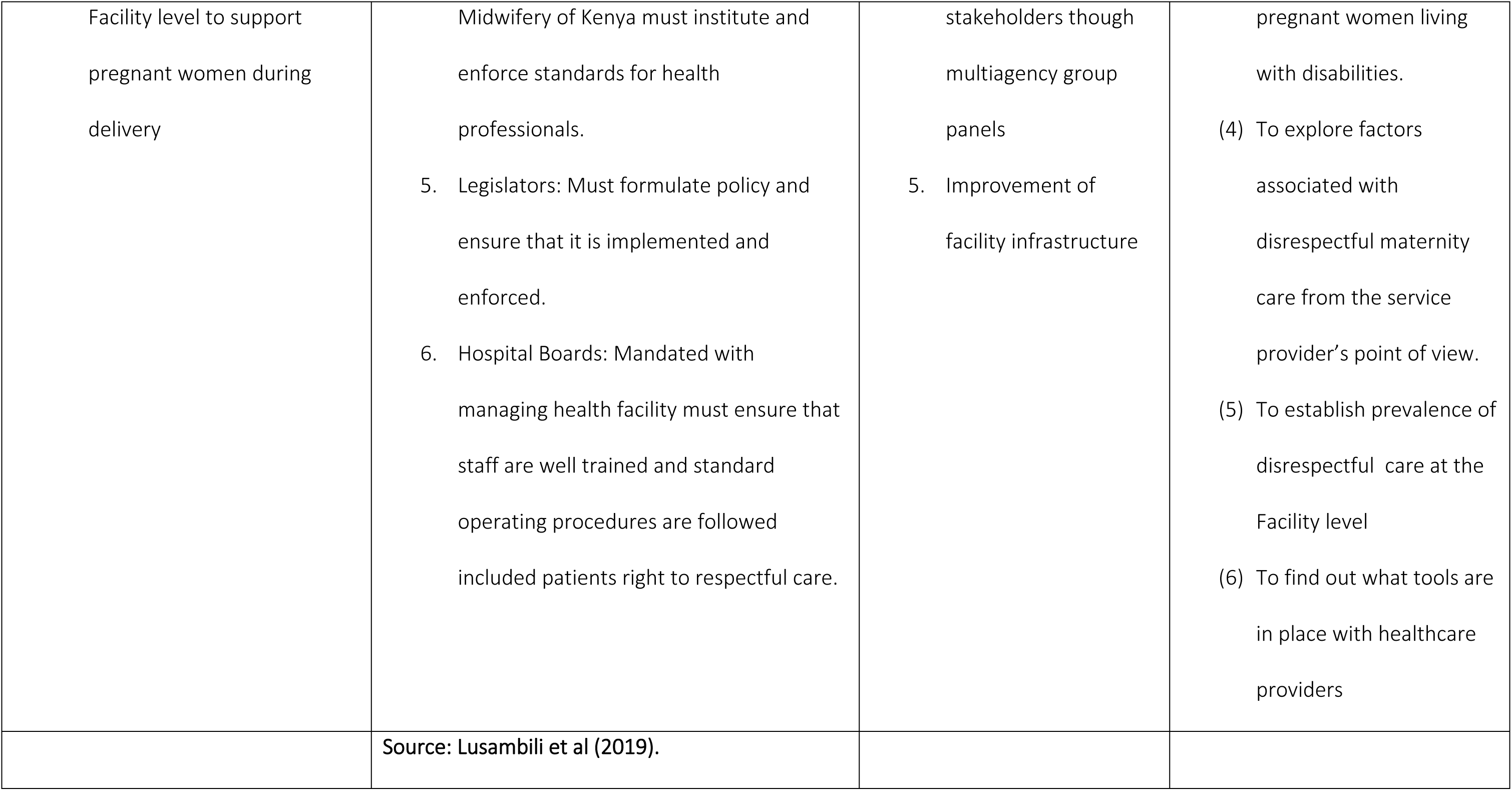
below provides a summary of our findings, suggestions and possible future research.

## CONCLUSIONS

This paper presents women’s experiences of disrespectful care during pregnancy, labour and delivery. Findings established that the health care service industry’s culture of disrespect and abuse has established a hostile environment for women in two communities’ maternity health care facilities that discourage their use for childbirth in future. And there is no reason to believe that these findings are not generalizable across other Kenyan communities. It is therefore not only bad for maternal and neonatal outcomes but also a significant barrier to the utilization of facility-based pregnancy and utilization throughout Kenya. A concerted effort from relevant stakeholders is needed to develop policies, standards, and intervention tools that can ensure gender responsive and respectful care for all women during pregnancy, labour and delivery. Stakeholders in various parts of the country including Ministry of Health (MOH), Kenya Medical Association (KMA), Kenya Nursing Association (KNA), and board members in different healthcare facilities have a vested interest and must all take steps to rectify disrespectful attitudes and practices that currently permeate Kenya’s public maternal health care services. Innovative approaches that are contextually congruent must be developed to integrate respectful maternity care as a routine quality component along a woman’s journey of pregnancy and delivery.

## STUDY STRENGTH

As noted in the methodology, the study elicited views from service users [women] and community members in two different contexts [men and local leaders] as well as interviews with key informants. This approach strengthened the data quality and trustworthiness meaning that the findings can be used to develop intervention tools in a rural context. The findings for this study are not intended to be generalizable in terms of statistical significance, but provide insight into challenges with promoting facility ante natal care, delivery and pre - natal care

## ACKNOWLEGMENTS

Authors wish to acknowledge all the study participants from the Kisii and Kilifi Counties. We acknowledge the funder –Government of Canada and Aga Khan Foundation Canada. We also acknowledge the feedback provided on the original manuscripts by Dr Abdulrahman Mohiddin, Sofia Jadavji and Dr Anisa Omar from Kilifi County.

## AUTHORS CONTRIBUTION

All authors have substantially contributed to the writing of this paper and have approved the version to published.

## DATA AVAILABILITY

Data cannot be shared publicly because of ethical consideration. Data are available from the Aga Khan University Monitoring and Evaluation Research Unit (MERL). Contact Institutional Data Access for researchers who meet the criteria for access to confidential data.

## REFERENCES

1. Gichuhi E, Lusambili A. iMedPub Journals Efficacy of Free Maternity Health Policy at Machakos Level 5 County Hospital (Kenya): An Exploratory Qualitative Study Keywords Beneficial changes observed after introduction. 2019;4–7.

2. Gitobu CM, Gichangi PB, Mwanda WO. The effect of Kenya’s free maternal health care policy on the utilization of health facility delivery services and maternal and neonatal mortality in public health facilities. BMC Pregnancy Childbirth. 2018;

3. Tunc Ö, Were W, Maclennan C, Oladapo O, Bahl R, Daelmans B, et al. Quality of care for pregnant women and newborns—the WHO vision. BJOG. 2015;

4. WHO. WHO recommendation on respectful maternity care during labour and childbirth. 2018;(February):1–11.

5. Alliance WR. Respectful maternity care: the universal rights of childbearing women. Survey Report. 2011.

6. Moindi RO, Ngari MM, Nyambati VCS, Mbakaya C. Why mothers still deliver at home: Understanding factors associated with home deliveries and cultural practices in rural coastal Kenya, a cross-section study Global health. BMC Public Health. 2016;

7. Bohren MA, Vogel JP, Tunçalp Ö, Fawole B, Titiloye MA, Olutayo AO, et al. Mistreatment of women during childbirth in Abuja, Nigeria: A qualitative study on perceptions and experiences of women and healthcare providers Prof. Suellen Miller. Reprod Health. 2017;

8. Kumbani L, Bjune G, Chirwa E, Malata A, Odland JØ. Why some women fail to give birth at health facilities: A qualitative study of women’s perceptions of perinatal care from rural Southern Malawi. Reprod Health. 2013;

9. Oyerinde K, Amara P, Harding Y. Barriers to Uptake of Emergency Obstetric and Newborn Care Services in Sierra Leone: A Qualitative Study. J Community Med Health Educ. 2014;

10. Namasivayam A, Osuorah DC, Syed R, Antai D. The role of gender inequities in women’s access to reproductive health care: A population-level study of Namibia, Kenya, Nepal, and India. Int J Womens Health. 2012;

11. Singh K, Bloom S, Haney E, Olorunsaiye C, Brodish P. Gender equality and childbirth in a health facility: Nigeria and MDG5. Afr J Reprod Health. 2012;

12. Banda PC, Odimegwu CO, Ntoimo LFC, Muchiri E. Women at risk: Gender inequality and maternal health. Women Heal. 2017;

13. Mmari K, Blum RW, Atnafou R, Chilet E, de Meyer S, El-Gibaly O, et al. Exploration of Gender Norms and Socialization Among Early Adolescents: The Use of Qualitative Methods for the Global Early Adolescent Study. Journal of Adolescent Health. 2017.

14. World Health Organization. WHO statement: The prevention and elimination of disrespect and abuse during facility-based childbirth. WHO press. 2015.

15. Mohale H, Sweet L, Graham K. Maternity health care: The experiences of Sub-Saharan African women in Sub-Saharan Africa and Australia. Women and Birth. 2017;

16. Gebrehiwot T, Goicolea I, Edin K, Sebastian MS. Making pragmatic choices: Women’s experiences of delivery care in Northern Ethiopia. BMC Pregnancy Childbirth. 2012;

17. Gebremichael MW, Worku A, Medhanyie AA, Edin K, Berhane Y. Women suffer more from disrespectful and abusive care than from the labour pain itself: A qualitative study from Women’s perspective. BMC Pregnancy and Childbirth. 2018.

18. Abuya T, Warren CE, Miller N, Njuki R, Ndwiga C, Maranga A, et al. Exploring the prevalence of disrespect and abuse during childbirth in Kenya. PLoS One. 2015;

19. Okafor II, Ugwu EO, Obi SN. Disrespect and abuse during facility-based childbirth in a low-income country. In: International Journal of Gynecology and Obstetrics. 2015.

20. Gabrysch S, Campbell OMR. Still too far to walk: Literature review of the determinants of delivery service use. BMC Pregnancy Childbirth. 2009;

21. Kyei NNA, Campbell OMR, Gabrysch S. The Influence of Distance and Level of Service Provision on Antenatal Care Use in Rural Zambia. PLoS One. 2012;

22. Tong A, Flemming K, McInnes E, Oliver S, Craig J. Enhancing transparency in reporting the synthesis of qualitative research: ENTREQ. BMC Med Res Methodol. 2012;

